# Functional divergence in a multi-gene family is a key evolutionary innovation for anaerobic growth in *Saccharomyces cerevisiae*

**DOI:** 10.1101/2022.06.15.496314

**Authors:** David J. Krause, Chris Todd Hittinger

**Affiliations:** Laboratory of Genetics, Wisconsin Energy Institute, DOE Great Lakes Bioenergy Research Center, Center for Genomic Science Innovation, J. F. Crow Institute for the Study of Evolution, University of Wisconsin-Madison, Madison, Wisconsin, USA

## Abstract

The amplification and diversification of genes into large multi-gene families often marks key evolutionary innovations, but this process often creates genetic redundancy that hinders functional investigations. When the model budding yeast *Saccharomyces cerevisiae* transitions from aerobic to anaerobic growth conditions, the cell massively induces the expression of seven cell wall mannoproteins (anCWMPs): *TIP1, TIR1, TIR2, TIR3, TIR4, DAN1*, and *DAN4*. Here we show that these genes likely derive evolutionarily from a single ancestral anCWMP locus, which was duplicated and translocated to new genomic contexts several times both prior to and following the budding yeast whole genome duplication (WGD) event. Based on synteny and their phylogeny, we separate the anCWMPs into four gene subfamilies. To resolve prior inconclusive genetic investigations of these genes, we constructed a set of combinatorial deletion mutants to determine their contributions toward anaerobic growth in *S. cerevisiae*. We found that two genes, *TIR1* and *TIR3,* were together necessary and sufficient for the anCWMP contribution to anaerobic growth. Overexpressing either gene alone was insufficient for anaerobic growth, implying that they encode non-overlapping functional roles in the cell during anaerobic growth. We infer from the phylogeny of the anCWMP genes that these two important genes derive from an ancient duplication that predates the WGD event, whereas the *TIR1* subfamily experienced gene family amplification after the WGD event. Taken together, the genetic and molecular evidence suggest that one key anCWMP gene duplication event, several auxiliary gene duplication events, and functional divergence underpin the evolution of anaerobic growth in budding yeasts.

## Introduction

Gene duplication is a common evolutionary process. Gene duplication events initially create redundancy, which can resolve itself in several ways, and for which myriad models have been proposed to explain the mechanisms of duplication and the fates of the gene duplicates (Innan and Kondrashov, 2010). A deeper understanding of the prevalence and conditions that favor these mechanisms requires rigorous testing of their respective hypotheses in model organisms. The birth-and-death model is a well-studied example in which multi-gene families undergo amplifications, followed by functional divergence or pseudogenization and gene loss (Nei et al., 1997). For example, this model has been applied to animal olfactory receptors (ORs), which have frequently undergone duplications followed by pseudogenization or functional differentiation (Hughes et al., 2018; Niimura and Nei, 2003). Identifying functional differentiation in OR genes is complicated by difficulties in assessing specific OR gene function and the effect on sensory perception, although mapping genetic variation to sensory perception differences can provide some information of functional divergence (Trimmer et al., 2019). This model has also been applied to genes in vertebrate immune systems, including major histocompatibility complex (MHC) and immunoglobulin genes, which have undergone many duplication and pseudogenization events (Nei and Rooney, 2005).

The model budding yeast *S. cerevisiae* and its close relatives are a powerful model system for studying the fates of gene duplicates due to their genetic tractability and a whole genome duplication (WGD) event that occurred in its ancestors circa 100 Mya. This WGD was a result of allopolyploidization between ancestors of the *Kluyveromyces/Lachancea/Eremothecium* (KLE) and *Zygosaccharomyces/Torulaspora* (ZT) yeast lineages (Marcet-Houben and Gabaldón, 2015; Wolfe and Shields, 1997). Most gene duplicates were ultimately lost, but among those that have been retained, several cases of sub-functionalization and neofunctionalization have been described (Cliften et al., 2006; Dean et al., 2008; Hickman and Rusche, 2007; Hittinger and Carroll, 2007; Kuang et al., 2016). Gene family expansions beyond single duplications are common for gene families found in the subtelomeres of yeast chromosomes, such as the *MAL*tose utilization genes, seri*PAU*perin genes, and FLOcculation genes (Brown et al., 2010; Luo and van Vuuren, 2009; Van Mulders et al., 2009). Studies of the aryl-alcohol dehydrogenase family and hexose transporter family in yeast have shown the power of *S. cerevisiae* as a system for studying multi-gene families, especially those with a high degree of redundancy, by successive individual gene deletions (Delneri et al., 1999; Wieczorke et al., 1999).

The WGD event also approximately coincides with the rise of anaerobic growth in this lineage of budding yeasts. While this trait is likely underpinned by several adaptations (Hagman et al., 2013; Thompson et al., 2013), one important adaptation is sterol transport (Ishtar Snoek and Yde Steensma, 2007; Snoek and Steensma, 2006). Sterol biosynthesis requires molecular oxygen, so sterols are not produced during anaerobic growth and are instead transported into the cell via the anaerobically-induced sterol transporters Pdr11 and Aus1 (Papay et al., 2020; Wilcox et al., 2002). Several studies have suggested that anaerobically-induced cell wall mannoproteins (anCWMPs) also play a role in this process (Alimardani et al., 2004; Inukai et al., 2015). Cell wall mannoproteins constitute nearly half of the fungal cell wall, along with β-glucans and chitin (Lipke and Ovalle, 1998). *S. cerevisiae* contains seven of these anCWMP genes, but the inference of their roles in anaerobic growth has been complicated by conflicting and inconclusive results. One study found that *tir3Δ, tir1Δ,* and *tir4Δ* single-mutant strains failed to grow anaerobically (Abramova et al., 2001), while another study found that a *tir1Δ* strain and a triple-mutant *tir1Δ tir2Δ tip1Δ* strain grew anaerobically without defect (Donzeau et al., 1996). Further complicating matters, three separate deletion library screens failed to identify any anCWMP deletion as having a detectable effect on anaerobic growth (Galardini et al., 2019; Reiner et al., 2006; Snoek and Steensma, 2006). In *Candida glabrata,* the *TIR3* homolog is required for anaerobic growth, but a *C. glabrata tir3Δ* mutant strain was not complemented by the *S. cerevisiae* homolog (Inukai et al., 2015). These conflicting results preclude the assignment of a definitive function for the anCWMP genes. Previous phylogenomic analyses found that the *DAN/TIR/TIP* family encoding anCWMPs underwent several gene duplication and translocation events after the WGD event in the lineage leading to *S. cerevisiae,* but the limited number of genomes available at the time hindered synteny analyses (Gordon et al., 2009). Thus, the family of genes encoding anCWMPs is ripe for both functional and evolutionary investigation.

Here we test a set of combinatorial anCWMP gene deletion mutants for growth in anaerobic conditions and find that only two genes are required for anaerobic growth, *TIR1* and *TIR3.* Together, these two genes are also sufficient for the anCWMP contribution to anaerobic growth. However, neither gene alone supports anaerobic growth, even when overexpressed, implying they encode distinct functional roles. We construct a phylogenetic tree of the all the anCWMP homologs in all published budding yeast genome sequences and show that this gene family is conserved across many taxa but underwent several gene duplication and translocation events around the time of the WGD. We infer that the *TIR1* and *TIR3* genes diverged prior to the WGD event in the family Saccharomycetaceae, which includes *S. cerevisiae,* and we propose hypotheses for the mechanism and timing of functional divergence and its relationship to subsequent gene duplication events and the evolution of anaerobic growth.

## Results and Discussion

### Contributions of the multi-gene anCWMP family to anaerobic growth

The anCWMP genes encode proteins that localize to the cell wall and have an extensively glycosylated serine/threonine rich domain. Despite genetic studies that have suggested some anCWMP genes may be involved in facilitating sterol transport (Alimardani et al., 2004; Inukai et al., 2015), a key feature of anaerobic growth, the genes still lack a clear biochemical or genetic function. We sought to definitively resolve conflicting prior genetic studies by constructing combinatorial deletion mutants in *S. cerevisiae.* Since *DAN1* and *DAN4,* as well as *TIR2* and *TIR4,* are adjacent in the genome, we deleted these two pairs together, resulting in five separate anCWMP loci deletions and 32 total combinations. We measured the growth of each resulting strain under anaerobic conditions (Fig. 1, Fig. S1).

**Figure 1.**
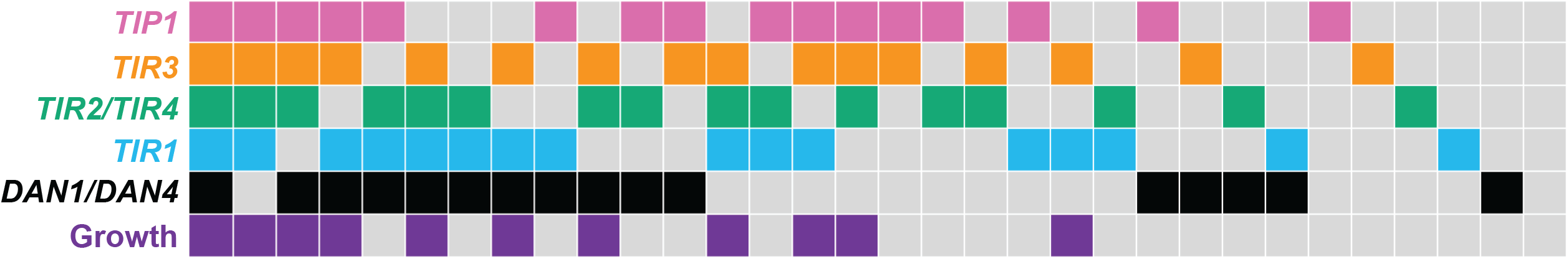
Growth of combinatorial mutant strains under anaerobic conditions. All 32 combinatorial mutant strains are represented by a single column, with presence of the anCWMP gene(s) depicted as a colored box and absence depicted by a gray box. The bottom row is shaded purple if the strain grew anaerobically to a final OD of greater than 0.5. Representative growth curves are shown in Figure SI.

Consistent with some previous studies, we found that *TIR3* was required for anaerobic growth (Abramova et al., 2001). No other anCWMP gene was essential for anaerobic growth. By limiting further analysis to strains containing *TIR3,* we found that all *TIR3*-containing backgrounds that failed to grow anaerobically were missing *TIR1.* The only strains lacking *TIR1* that could grow anaerobically contained *TIR3, TIR2/TIR4,* and either *TIP1* or *DAN1/DAN4.* Further, the presence of *TIR1* and *TIR3* was sufficient to yield anaerobic growth in the absence of the other five anCWMP genes. We infer that *TIR1* contributes to anaerobic growth, but this contribution may be partially redundant with other anCWMP genes. The unknown anCWMP requirements for anaerobic growth have hampered attempts to fully engineer sterol transport into yeasts that naturally lack sterol uptake or into *S. cerevisiae* under aerobic conditions (Alimardani et al., 2004). We hypothesize that *TIR1* and *TIR3* encode the minimal anCWMP functions necessary for anaerobic growth and sterol uptake.

Why then does *S. cerevisiae* contain seven different anCWMP genes if two are sufficient for anaerobic growth? The positive dosage model of gene duplication describes a situation in which gene duplicates are retained because the resulting increased expression of the gene pair provides a fitness benefit (Kondrashov et al., 2002). We reasoned that the presence of seven anCWMP genes in the *S. cerevisiae* genome might be the result of benefits from increased gene expression of this set of genes with potentially overlapping functions. We tested this hypothesis by expressing each anCWMP gene individually on a high-copy number plasmid in the seven-gene deletion mutant. No single anCWMP gene conferred anaerobic growth to this seven-gene deletion mutant (Fig. 2A). Given that no single locus conferred anaerobic growth even when its dosage was artificially increased via a high-copy plasmid, we reject the positive dosage model as a sufficient explanation for the maintenance of the seven anCWMP genes.

**Figure 2.**
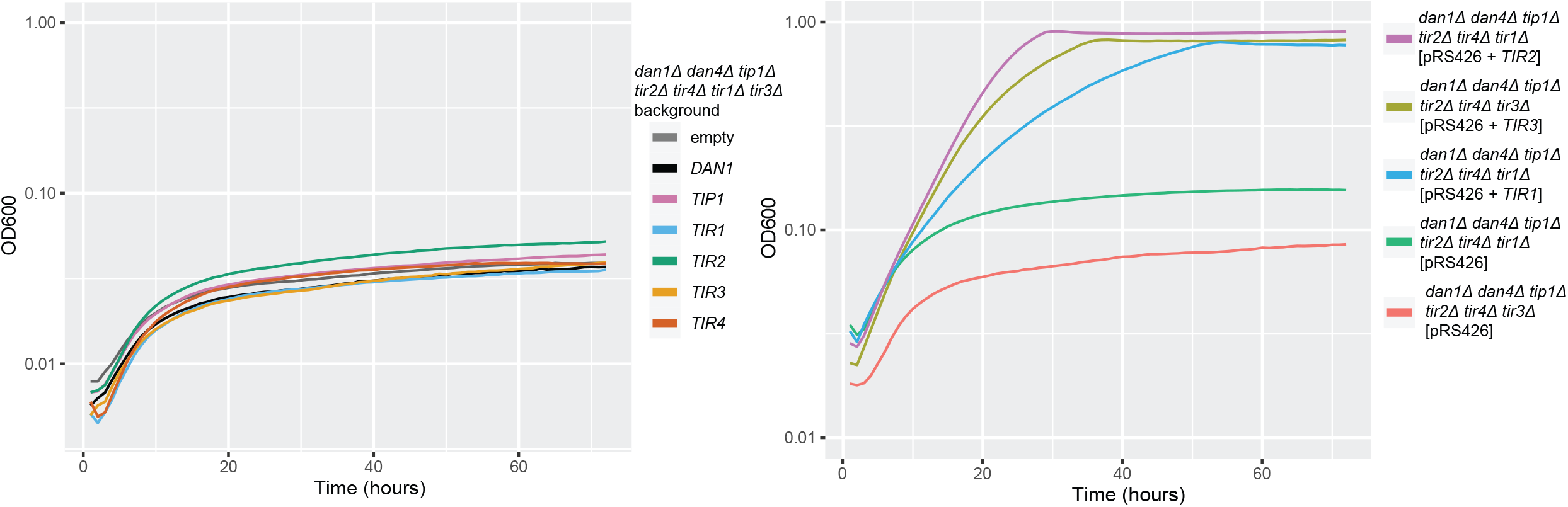
Growth of strains carrying anCWMP genes on a 2μ high-copy vector with log-scaled Y-axes. A) Representative growth curves for the strain lacking all genomic anCWMP genes but carrying one anCWMP gene on the high-copy vector. B) Representative growth curves for backgrounds lacking all genomic anCWMP genes except either *TIR3* or *TIR1,* but carrying one anCWMP gene on the high-copy vector. All three strains that grew are shown here, but all tested strains are shown in Figure S2.

Since *TIR3* is required but not sufficient for anaerobic growth, either at its native locus or when overexpressed, we performed the same overexpression experiment in a strain containing only *TIR3* (i.e. the other six genes were deleted). As expected, overexpression of *TIR1* conferred anaerobic growth in the *TIR3*-containing background. We also found that overexpression of *TIR2* conferred growth, even though a *TIR3*-containing strain containing the *TIR2/TIR4* native locus did not grow (Fig. 2B, Fig. S2). This result implies that, when overexpressed, *TIR2* may complement the function of *TIR1,* but its native expression level is not high enough to normally do so. To test this hypothesis, we analyzed published RNA-sequencing data from *S. cerevisiae* under anaerobic conditions and found that *TIR2* gene expression was 50-fold lower than *TIR1* (Myers et al., 2019). This result is consistent with the combinatorial mutant experiment, in which *tir1Δ* mutant backgrounds that grew always contained *TIR3, TIR2/TIR4,* and another anCWMP gene. We also overexpressed some anCWMP genes in a strain containing only *TIR1* (i.e. the other six genes were deleted). In this case, overexpression of *TIR3* conferred anaerobic growth, while *TIP1* and *TIR2* did not (Fig. 2B, Fig. S2). We conclude that *TIR3* and *TIR1* encode distinct functions that are both required for anaerobic growth, but the *TIR1* function is partially redundant with other anCWMP genes.

### The anCWMP genes comprise at least four subfamilies

To investigate the evolution of this multi-gene family, we searched 332 publicly available budding yeast genomes (Shen et al., 2018) for homologs of the *S. cerevisiae* anCWMP genes, and we found homologs in most species. We also found homologs of the closely related gene *AFB1* in approximately 40% of species, and this gene family clustered outside the anCWMP genes with 100% bootstrap support (Fig. 3a). Due to the low complexity of the serine/threonine-rich region present in the middle of these gene sequences, only the short, structured N-terminal portion and the short glycosylphosphatidylinositol (GPI)-anchoring C-terminal portion were used for alignment and phylogenetic analyses (Fig. 3b). The seven members of the anCWMP gene family in *S. cerevisiae,* as well as all the anCWMP genes from the family Saccharomycetaceae, formed a single clade within the larger gene tree (Fig. 3a, Fig. S2). This result implies that a gene family expansion occurred within the Saccharomycetaceae, so we will focus here on that clade. Even within the family Saccharomycetaceae, most non-WGD species do not contain any anCWMP genes, but those that do contain them harbor between one and four genes in a single genomic neighborhood. This gene neighborhood is generally most similar to the *TIR1* gene neighborhood of *S. cerevisiae.* In the post-WGD lineage, all species except *Vanderwaltozyma polyspora* contain anCWMP genes, which number as few as three in *Candida (Nakaseomyces) castellii* and as many as 17 in *Kazachstania unispora.*

**Figure 3.**
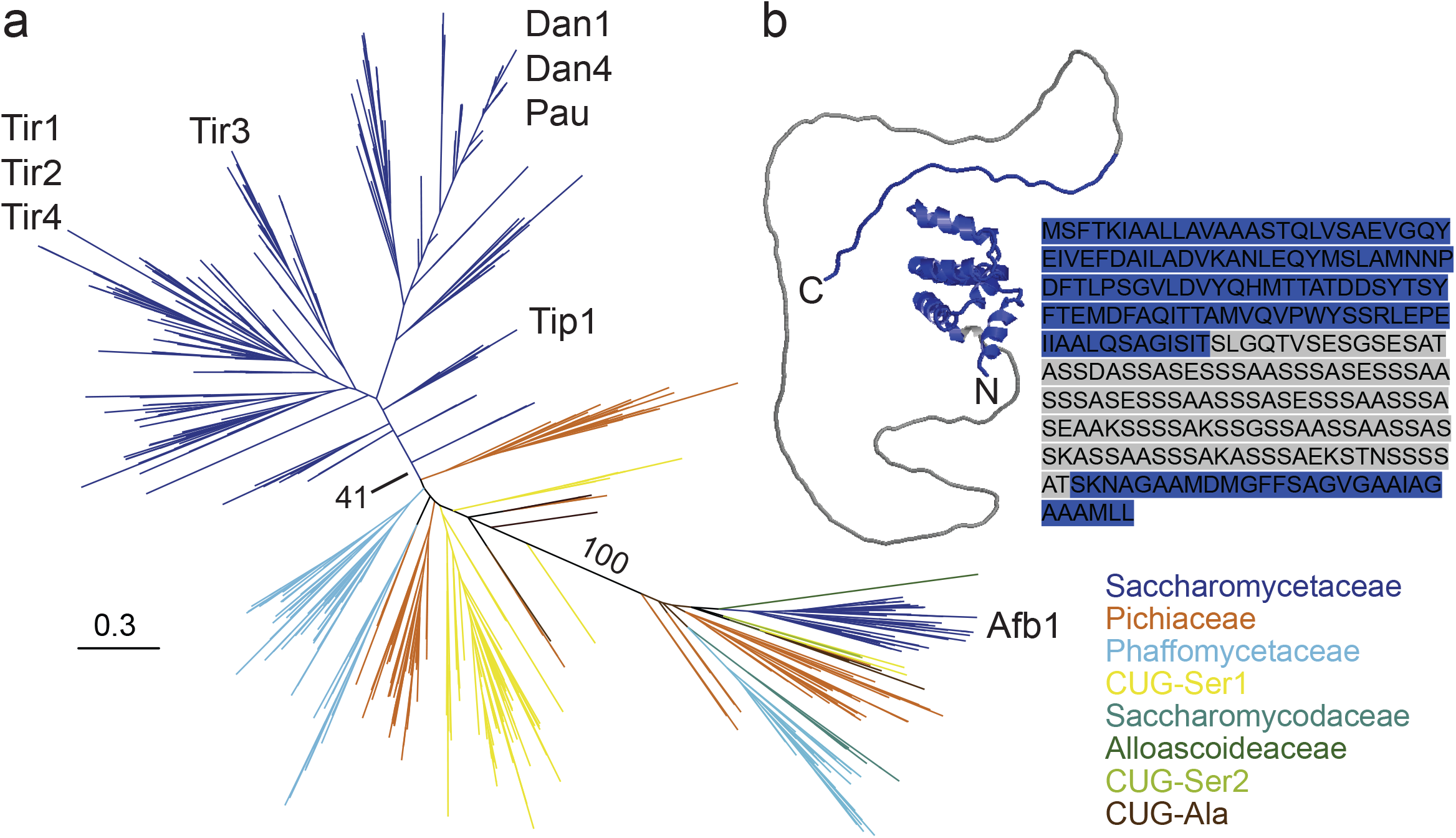
A) Maximum likelihood phylogeny of the anCWMP genes and *AFB1* genes from the budding yeast subphylum Saccharomycotina using amino acids. Yeast major clades are colored according to Shen et al. 2018. Key bootstrap values are shown at the origin of Saccharomycetaceae anCWMP genes and the clustering of anCWMP genes from *AFB1.* B) Predicted protein structure from alpha-fold along with amino acid sequence of *TIR3* from *S. cerevisiae* (Jumper et al., 2021; Varadi et al., 2022). Blue regions depict those used for alignment and phylogenetic analyses: the N-terminal structured region and the C- terminal GPI-anchor signal sequence, which is ultimately cleaved. The grey region depicts the serine/threonine-rich region, which was removed from protein sequences to facilitate alignments.

The phylogeny of the anCWMP genes revealed several distinct clades, which we designate here as four subfamilies of the anCWMP gene family based on their related *S. cerevsiae* homologs: a) *TIR3;* b) *TIR1*, which includes *TIR1, TIR2,* and *TIR4;* c) *TIP1;* and d) *DAN1*, which includes *DAN1* and *DAN4* (Fig. 3, Fig. 4). The subfamilies either have strong bootstrap support, shared synteny patterns, or both. We discuss each subfamily in more detail below, as well as those homologs that do not neatly fit into the subfamilies that we currently recognize.

**Figure 4.**
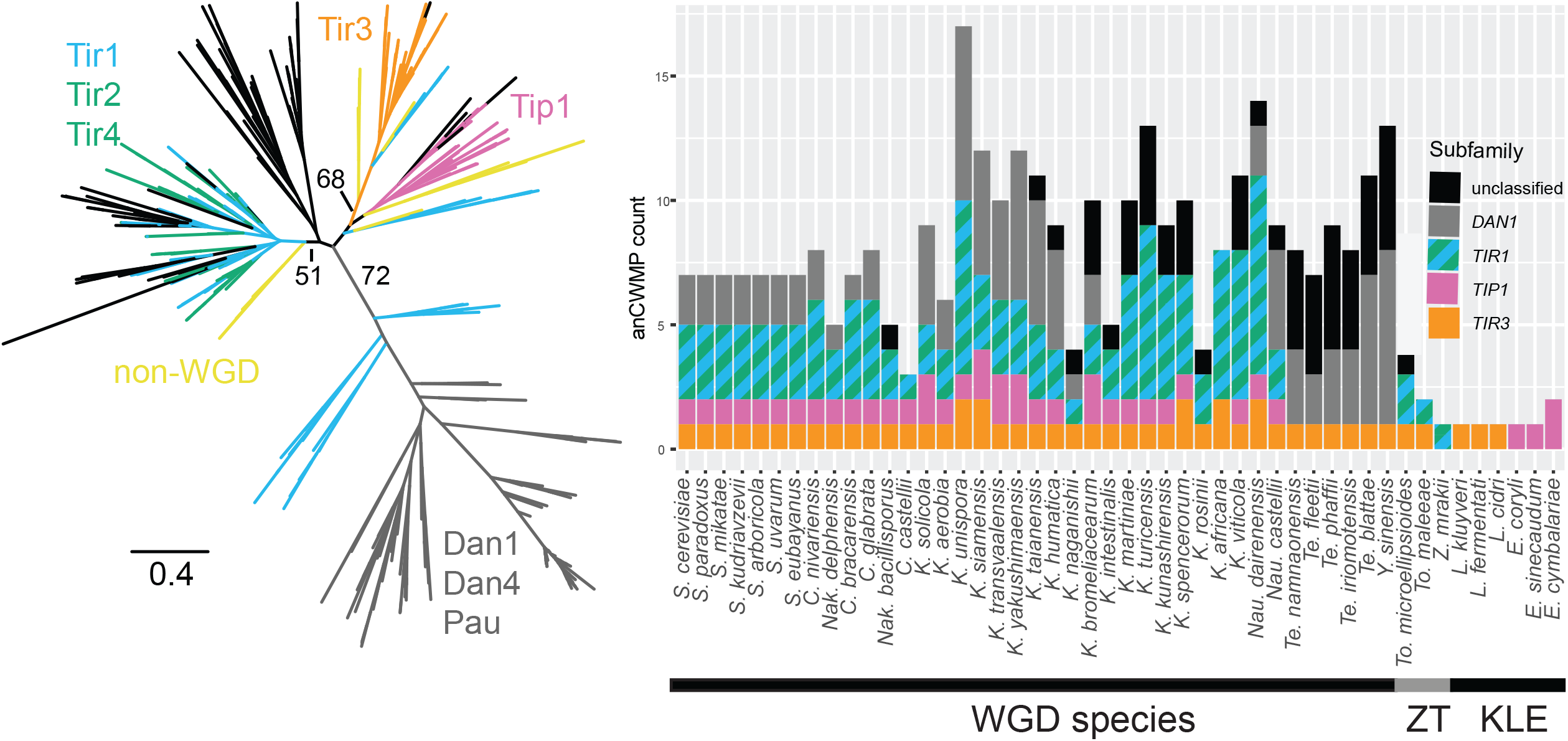
A) Maximum likelihood phylogeny of the anCWMP genes in Saccharomycetaceae using amino acid sequences. Branch values shown are the result of 100 bootstrap replicates. Branches are colored based on synteny with the *S. cerevisiae* homolog. Yellow branches indicate genes from non-WGD species. B) Total number of each inferred anCWMP subfamily in each species of Saccharomycetaceae containing homologs. Subfamilies were defined as clades with bootstrap support values greater than 50% or shared synteny of 90% of its members with the *S. cerevisiae* homolog.

Genes related to *TIR3* formed a clade with 68% bootstrap support, and this subfamily was the most widely conserved; indeed, all post-WGD species in the dataset and several non-WGD species contain a putative *TIR3* ortholog. The members of this subfamily were generally syntenic with *S. cerevisiae TIR3,* except for the homologs in *Tetrapisispora/Yueomyces* and non-WGD species, which were found in the putative ancestral locus of all anCWMP genes. This result implies that the structure of the *TIR3* locus in *S. cerevisiae* and its relatives resulted from a translocation event after divergence from the *Tetrapisispora/Yueomyces* lineage. The presence of a *TIR3* ortholog in both ZT clade members and KLE members implies that this gene was present in both parents of taxa descended from the WGD event. Because most post-WGD species contain only one copy of *TIR3* with a few exceptions resulting from recent duplications, one copy of the *TIR3* gene was likely lost sometime after the WGD event.

Genes related to *TIR1, TIR2,* and *TIR4* formed a clade with 51% bootstrap support, and all post-WGD species, except *Kazachstania transvaalensis* and *Tetrapisispora/Yueomyces,* contain at least one homolog. These genes generally shared synteny with the *TIR1, TIR2,* or *TIR4* genes of *S. cerevisiae,* but there was no clear phylogenetic distinction between genes sharing synteny with *TIR1* and those sharing synteny with the *TIR2/TIR4* locus. Instead, gene members of this subfamily were generally more closely related to other members within a particular species or small clade of species than they were to homologs in other species. Concerted evolution homogenizes several copies of a multi-gene subfamily via recombination, and this process may be adaptive when gene dosage is critical to function (Hurst and Smith, 1998). Indeed, the overlapping functions of *TIR1* and *TIR2* demonstrated in our genetic complementation assays and the redundancies in our combinatorial mutant analyses support this model of dosage-dependent homogenization. Several members of the *Torulaspora* and *Zygotorulaspora* genera have one or more members of the *TIR1* subfamily, and these originate at the base of the *TIR1* clade, suggesting that this subfamily was present in the ZT parent of the WGD hybridization (Fig. 4: yellow taxa at base of *TIR1* clade).

Genes related to *TIP1* formed a clade with 25% bootstrap support, and all post-WGD species, except *Kazachstania rosinii, Kazachstania africana, Tetrapisispora* spp., and *Y. sinensis,* contain at least one homolog. This subfamily may also include the anCWMP genes from *Eremothecium* spp., which branched at the base of the *TIP1* clade, were found in the ancestral locus of all anCWMP genes, and had a bootstrap support of 14%. Similar to the logic applied for *TIR3* and *TIR1,* this result is consistent with the presence of *TIP1* in the KLE parent of the WGD event, which was then translocated after the WGD. We infer that *TIP1* was subsequently lost in the *Tetrapisispora/Yueomyces* clade.

Genes related to *DAN1* and *DAN4* formed a clade with 72% bootstrap support. Their long branch lengths suggest that these genes rapidly diverged from the other anCWMP genes, and most post-WGD species contain at least one homolog, except some *Nakaseomyces* and *Kazachstania* species. Genes of this subfamily are generally located in the subtelomeres of chromosomes or near the ends of small contigs. *Tetrapisispora* spp. and *Yueomyces sinensis DAN1* genes are informative exceptions because they are located near the putative ancestral locus of all anCWMP genes (blue-colored clades at base of *DAN1* subfamily in Fig. 4). No non-WGD species contain clear *DAN1* homologs. These data imply that the *DAN1* gene subfamily emerged after the WGD event, and that subsequent translocation to subtelomeric regions of chromosomes did not occur in the *Tetrapisispora/Yueomyces* lineage. The *PAU* genes present in *Saccharomyces* spp. are embedded within this clade, and they share recent ancestry and subtelomeric localization with the *DAN* genes.

These classifications exclude a small set of well-supported clades that lacked clear relationships with *S. cerevisiae* subfamilies. These divergent homologs may represent additional subfamilies and include several clades of genes found in *Naumovozyma* spp., *Kazachstania* spp., and *Tetrapisispora/Yueomyces* spp. Due to the poor bootstrap values for these clades and their phylogenetic placement basal to well-supported subfamilies, it is unclear whether these are divergent homologs of recognized subfamilies or additional novel subfamilies.

### Origins of the anCWMP genes, their divergence, and their role in anaerobic growth

To better understand the evolution of the anCWMP gene family and their function, we further investigated this gene family in the context of all budding yeast species and its relationship to the homolog *AFB1*. We found several homologs of the *AFB1* gene as significant hits while searching for homologs of the anCWMP gene family in budding yeasts. We did not find homologs of either *AFB1* or the anCWMP genes in species outside the budding yeast subphylum, nor in any of the more basal budding yeast major clades, such as Lipomycetaceae, Trigonopsidaceae, or Dipodascaceae/Trichomonascaceae. *AFB1* encodes an a-factor barrier protein that is believed to bind to a-factor that is secreted from *MAT***a** cells, but it is not required for mating (Huberman and Murray, 2013). The anCWMP genes and *AFB1* share sequence similarity in their N-terminal structured regions, as well as a low-complexity serine/threonine-rich region and a putative GPI-anchoring C-terminal region. The precise role of the anCWMP genes during anaerobic growth may be related to the putative function of Afb1, which binds in the cell wall to isoprenoid-related molecules that are structurally similar sterols.

While *AFB1* is predominantly a single-copy gene when present in budding yeasts, the anCWMP genes are often multi-copy, with 46% of species that contain anCWMP genes possessing two or more. This multi-copy nature is likely indicative of lineage-specific amplifications: copy number varies widely between yeast clades, while anCWMP genes within yeast clades tend to cluster together, rather than with homologs from other yeast clades. Although six species within the Phaffomycetaceae, *Babjeviella inositovorans,* and the *Brettanomyces* spp. all contain four or more anCWMP genes, here we focused on the most striking gene family amplification that occurred within the post-WGD lineage of Saccharomycetaceae. Functional characterization of the anCWMP genes outside of *S. cerevisiae* will better illuminate the roles of these genes in budding yeasts. The amplifications within Saccharomycetaceae and *Brettanomyces* are particularly interesting because they coincide with the independent evolution of these clades’ abilities to grow anaerobically (Visser et al., 1990).

Our genetic experiments in *S. cerevisiae* show that the *TIR1* and *TIR3* genes are the major cell wall mannoprotein contributors to anaerobic growth. Establishing the precise timing of divergence between these two genes is challenging due to the general difficulty in determining the relationships between subfamilies using the few alignable sites. Bootstrap supports were low on branches connecting subfamilies (Fig. 4a), and phylogenetic topology tests failed to reject any tree topologies (Table S1). We attempted to root the Saccharomycetaceae subtree using anCWMP genes from Phaffomycetaceae or *AFB1* genes. Much like our attempts to determine relationships between subfamilies, the two trees were inconsistent in root placement and tree topology, demonstrating the difficulty in reliably assessing the evolutionary relationships among the anCWMP subfamilies (Fig. S4). Nonetheless, we can conclude that the anCWMP genes in the Saccharomycetaceae arose via several duplication and divergence events.

We can also infer that the *TIR1* and *TIR3* genes diverged prior to the WGD event because the non-WGD species *Torulaspora microellipsioides* and *Torulaspora maleeae* both contain putative members of both subfamilies. This result implies that the ZT parent of the WGD allopolyploidization event contained both genes. Given the presence of *TIR3* homologs in multiple *Lachancea* spp., the KLE parent likely contained at least a *TIR3* gene as well. Future work will be needed to determine whether the non-WGD homologs of these genes can functionally replace their post-WGD counterparts and to contribute to our understanding of the timing of the functional divergence between *TIR1* and *TIR3.* The timing of critical functional divergence is also complicated by the absence of clear *TIR1* homologs in the *Tetrapisispora/Yueomyces* lineage, a majority of whose species we found to grow under anaerobic conditions (Table S2). Future work could also address whether other anCWMP genes have evolved to functionally replace *TIR1* function in these species.

The anCWMP genes present an interesting system for studying the birth-and-death model of evolution, in which multi-gene families experience recurring duplications followed by functional divergence, dosage changes, or gene loss events. While we focused here on *S. cerevisiae* and its close relatives, future genetic experiments in *Brettanomyces* under anaerobic growth conditions, as well as experiments to identify functions of the expanded gene family members in other yeast lineages, would further contribute to understanding the birth-and-death process, as well as the cryptic functions of these genes. One prime target for future study is the genus *Kazachstania,* where every anCWMP gene subfamily has experienced either lineage-specific amplification events, gene loss events, or both. The *PAU* genes of the genus *Saccharomyces* are another example of genes that have experienced a lineage-specific amplification within the genus followed by many loss events, and functional characterization of these genes may be on the horizon with genetic tools, such as CRISPR-Cas9. Further work in diverse budding yeast species will continue to shed light on how duplication, functional divergence, and gene loss in the CMWP gene family have occurred in various yeast lineages and what phenotypic effects these processes have had.

Here, we have identified a minimal set of two anCWMP genes, *TIR3* and *TIR1,* that are necessary and sufficient for anaerobic growth, likely by supporting sterol transport in *S. cerevisiae.* This finding may facilitate engineering sterol uptake into naïve yeast species or into *S. cerevisiae* under conditions when sterol uptake is normally repressed. This minimal set of genes provides a simplified system for studying the function of cell wall mannoproteins, as well as the functional divergence that underlies their mutual necessity for anaerobic growth. While we identified a critical anaerobic role for these genes, understanding their function in obligate aerobic species will yield insights into the evolutionary origins of anaerobic growth. The contemporaneity of anCWMP gene family expansions and origins of anaerobic growth that have independently occurred within the Saccharomycetaceae and the distantly related genus *Brettanomyces* suggest that gene amplification may be a critical step in the evolution of anaerobic growth. The phylogenetic relationships and functional differentiation among *S. cerevisiae* anCWMP genes observed here for the first time shed vital light on the origins of this ecologically and industrially important trait and set the stage for broader investigations.

## Materials and Methods

### Strains, media, and oligonucleotides

All genetic manipulations were performed in the prototrophic *S. cerevisiae* S288C *MATα* strain *(SUC2 gal2 mal2 mel flo1 flo8-1 hap1 ho bio1 bio6*). Gene replacement mutants using the *kanMX, hygMX, natMX,* and *zeoMX* cassettes were selected on 200mg/L G418 (US Biological Life Sciences), 300mg/L Hygromycin B (US Biological Life Sciences), 100mg/L ClonNAT (WERNERBioAgentsGmbH), and 100mg/L Zeocin (Invitrogen), respectively. The *tip1-Δ* markerless deletion was obtained by first replacing the *TIP1* CDS with the *URA3* marker, then counterselecting against the *URA3* marker with a markerless repair template. All yeast strains used in this study can be found in Table S3 Oligonucleotide sequences for construction and screening of the mutants can be found in Table S4.

Strains were generally grown in synthetic complete (SC) consisting of 5g/L ammonium sulfate, 1.72g/L yeast nitrogen base, 2g/L synthetic dropout mix, and 20g/L glucose (all reagents from US Biological Life Sciences). SC without uracil was used for strains carrying *URA3-se*lection plasmid, and the synthetic dropout mix for this medium lacked uracil. For counterselection against *URA3,* SC plates containing 1g/L 5’-FOA and 50mg/L additional uracil were used (US Biological Life Sciences). YPD medium contained 10g/L yeast extract, 20g/L peptone, and 20g/L glucose.

### Backcrossing and collecting haploids

The *MATα dan1-dan4Δ tir2-tir4Δ tir3Δ tir1Δ tip1Δ ura3Δ* strain and a *MATa met6Δ* strain were spotted together and grown overnight on a minimal medium plate (lacking methionine and uracil). Colonies that grew were restreaked to a minimal medium plate and confirmed by PCR to be diploid at the *MAT* locus. The diploid was then streaked to a GNA presporulation plate (50g/L glucose, 30g/L Difco nutrient broth, 10g/L yeast extract, 20g/L agar) overnight, followed by inoculation of cells to sporulation medium for 72 hours at room temperature. The mix of spores and unsporulated diploids was then centrifuged, and the pellet was incubated with yeast protein extraction reagent (Thermo Scientific) and vortexed, followed by several washes with sterile dH_2_O. The cells were then plated to YPD, and colonies were recovered and screened by PCR at the *MAT* locus and all anCWMP loci to confirm ploidy and determine genotypes.

### Anaerobic growth experiments

*A* portion of a yeast colony was picked from a YPD plate into YPD medium for overnight growth at 30°C. The next morning, the saturated culture was diluted 20-fold into SC medium and grown for four hours at 30°C. These actively growing cultures were then introduced into a Coy anaerobic chamber and diluted 50-fold into an anaerobic 96-well plate containing 200μl of anaerobic medium and placed on a Tecan Spark-Stacker. The plate reader read absorbance or optical density (OD) at 600nm once every hour after 5s of shaking. After 24 hours of anaerobic culturing, the cultures were diluted 50-fold for a second round of growth, and these data form the results of the anaerobic experiments. For strains carrying plasmids with *URA3* selection, all YPD and SC media were replaced with SC medium lacking uracil. All anaerobic media contained anaerobic supplements of 20μg/mL ergosterol and Tween80.

### Generating the anCWMP gene sequences within Saccharomycotina

The amino acid sequences of all seven anaerobic CMWP genes from *S. cerevisiae* were used to search the publicly available genome sequences of 332 yeast species via http://y1000plus.org/blast using an evalue cutoff of 0.01 (Priyam et al., 2019; Shen et al., 2016). Full gene sequences were manually extracted from the genome sequence files using coordinates from the blast outputs. Genes containing ‘N’ nucleotide calls due to scaffolding or sequencing errors and multi-domain anCWMP genes were excluded from the gene list. The full-length sequences used in this study can be found in Supplementary File 1. Neighboring genes in *S. cerevisiae* were used as queries to identify homologs in the target genomes, and a anCWMP gene was considered syntenic with the *S. cerevisiae* homolog if a shared neighboring gene was found within 10Kb of the anCWMP homolog (Table S5). Because of alignment difficulties presented by the serine/threonine-rich region, this region was removed by concatenating the N-terminal portion to the last thirty codons encoding the C-terminal portion (see Fig. 3B, sequences in Supplementary File 2). Amino acid sequences were aligned using MAFFT (Katoh and Standley, 2013), and alignment positions with greater than 25% gaps were removed using Trimal version 3 (Capella-Gutiérrez et al., 2009). Maximum-likelihood phylogenies were constructed using RAxML v.8.2.11 with the PROTGAMMAAUTO parameter and 100 rapid bootstrap calculations(Stamatakis, 2014). AU tests were performed using IQTREE v.1.6.8 (Nguyen et al., 2015).

---

Figure S1 – Representative growth curves for each of the 32 strains depicted in Fig. 1. The curve for each strain was selected from three biological replicates and represents the median final saturation value.

Figure S2 – Representative growth curves for strains deleted for all anCWMP genes except *TIR1* or *TIR3*, with multi-copy plasmids carrying individual anCWMP genes.

Figure S3 – Maximum likelihood anCWMP gene tree constructed using Saccharomycetaceae homologs and rooted using either A) Phaffomycetaceae anCWMP gene homologs or B) *AFB1* homologs. Trees were built using aligned amino acid sequences.

Supplementary File 1 – Amino acid sequences with species names for anCWMP gene and *AFB1* homologs used in alignments and phylogenies in this study.

Supplementary File 2 – Amino acid sequences used in this study with serine/threonine-rich regions removed.

Supplementary File 3 – Newick tree file with taxon names for tree depicted in Fig. 3A.

Supplementary File 4 – Newick tree file with taxon names for tree depicted in Fig. 4A.

## Acknowledgments

We thank Robert A. Sclafani for S288C and Trey K. Sato, Kaitlin J. Fisher, and lab members for helpful suggestions. This material is based upon work supported in part by the Great Lakes Bioenergy Research Center, U.S. Department of Energy, Office of Science, Office of Biological and Environmental Research under Award Number DE-SC0018409; the National Science Foundation under Grant Nos. DEB-1442148 and DEB-2110403; and the USDA National Institute of Food and Agriculture (Hatch Project 1020204). C.T.H. is a H. I. Romnes Faculty Fellow, supported by the Vice Chancellor for Research and Graduate Education with funding from the Wisconsin Alumni Research Foundation.

## Data Availability

The data underlying this article are available in the article and in its online supplementary material.

