## Supplementary figures and images for "Functional divergence in a multi-gene family is a key evolutionary innovation for anaerobic growth in *Saccharomyces cerevisiae*"

### Figure S1

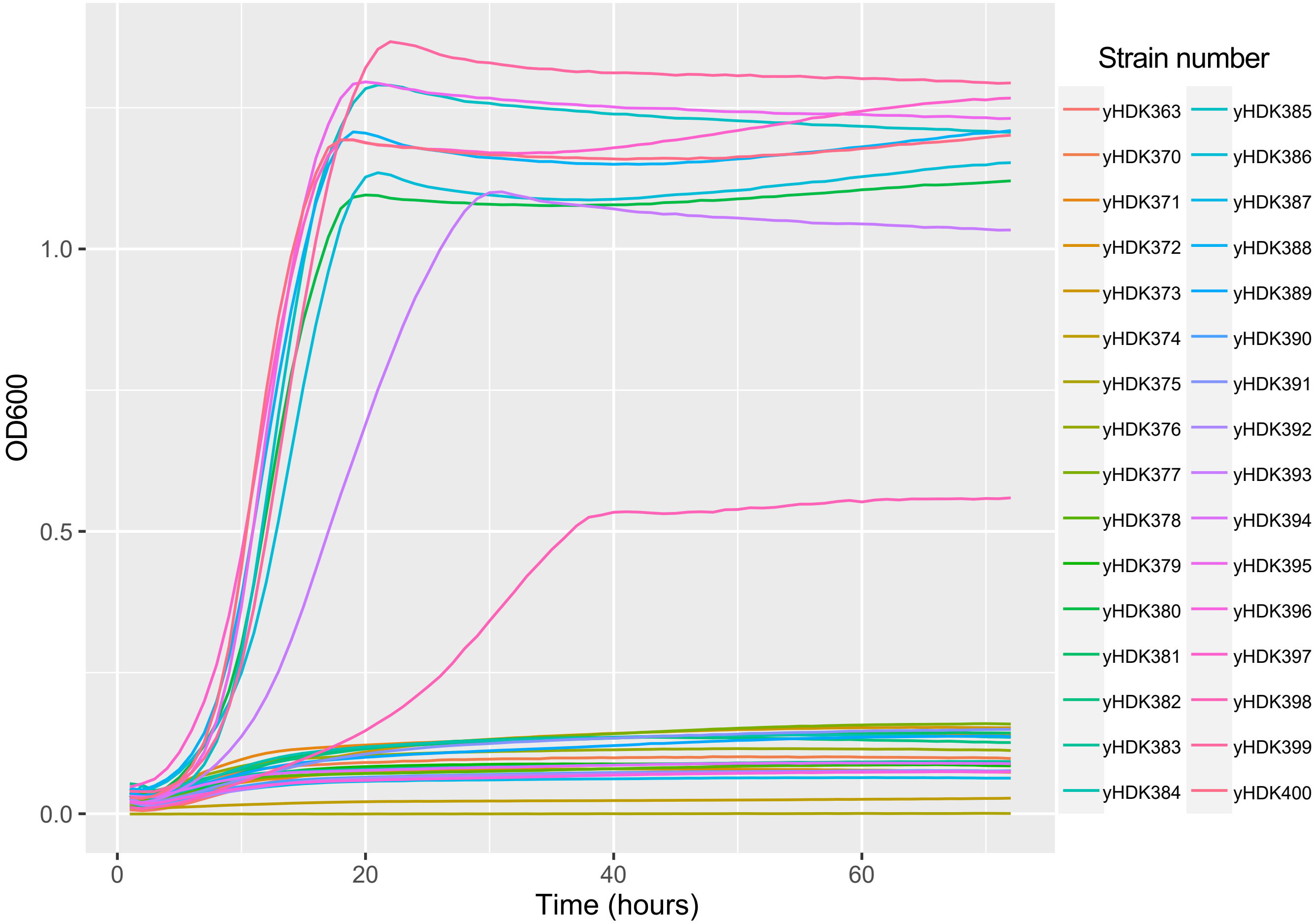

### Figure S2

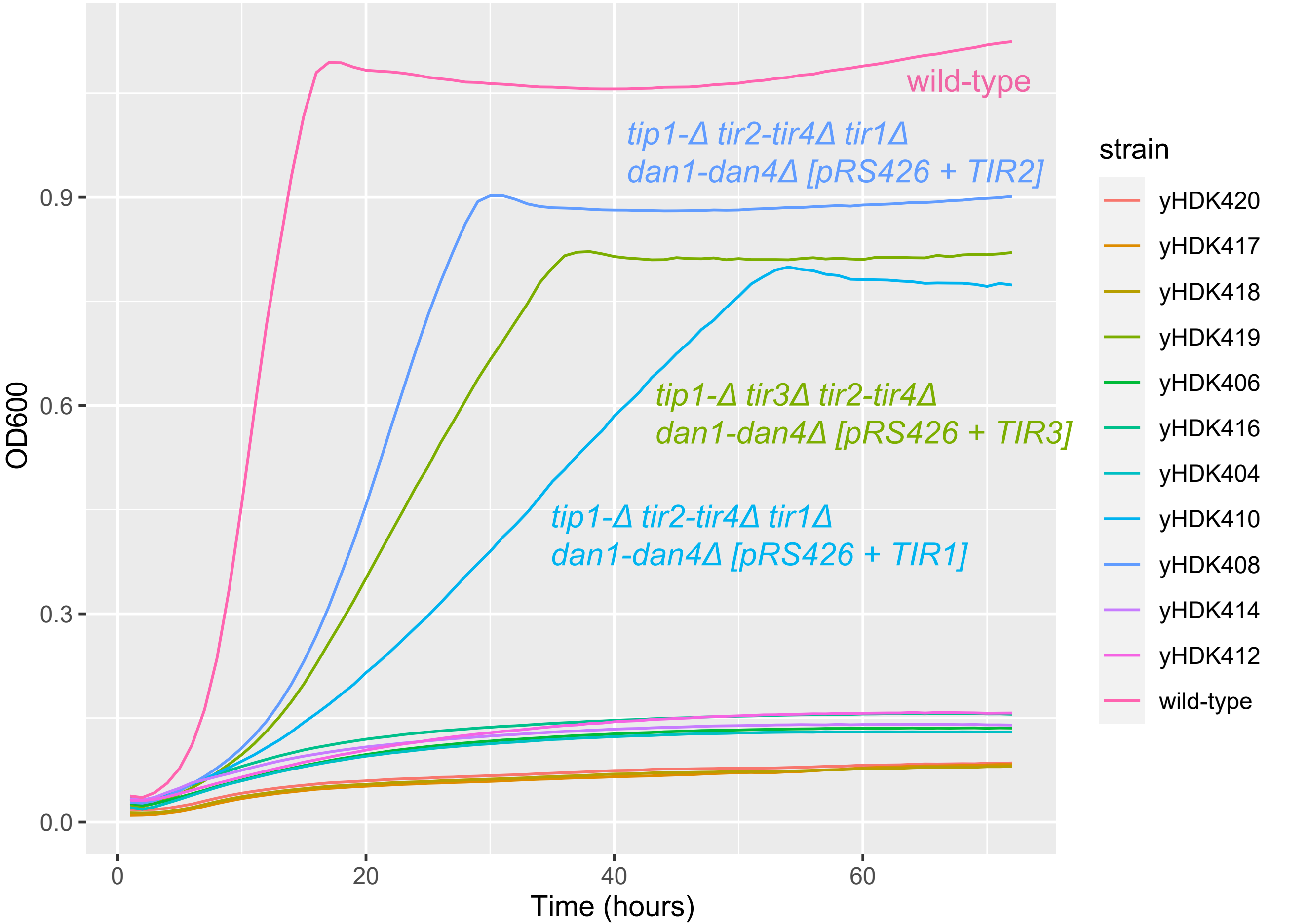

### Figure S3

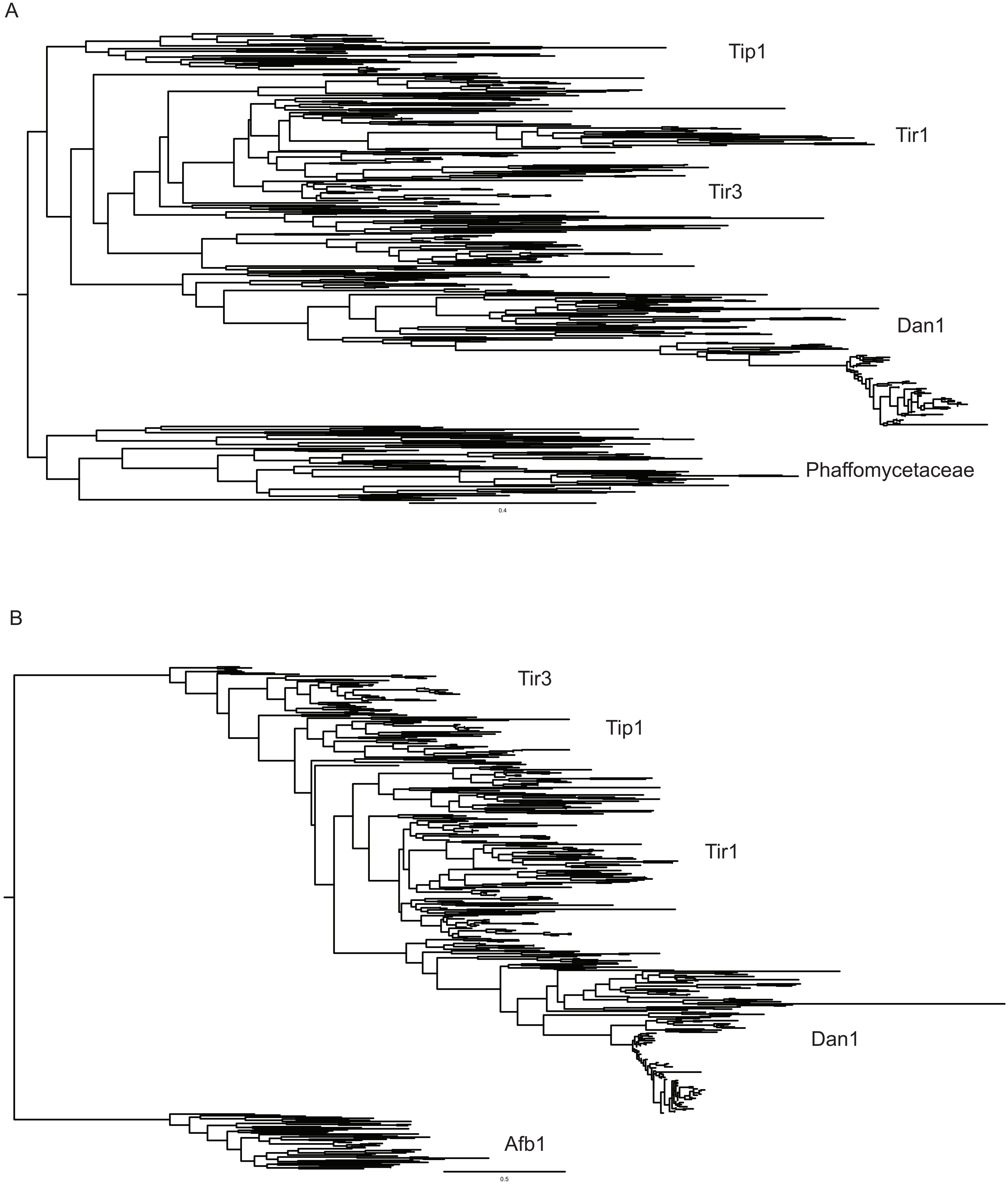
