## Supplementary material for "Functional divergence in a multi-gene family is a key evolutionary innovation for anaerobic growth in *Saccharomyces cerevisiae*": Table S2

Table S2 – Anaerobic growth at 48 hours after subculturing for selected species.

| Species | OD at 48 hours |
| --- | --- |
| *Saccharomyces mikatae*^a^ | 1.1 ± 0.032 |
| *Tetrapisispora iriomotensis* | 0.78 ± 0.12 |
| *Tetrapisispora fleetii* | 0.90 ± 0.010 |
| *Tetrapisispora namnaonensis* | 1.0 ± 0.039 |
| *Tetrapisispora phaffii* | 0.046 ± 0.00083 |
| *Dekkera anomala* | 0.76 ± 0.045 |
| *Kluyveromyces marxianus*^b^ | 0.10 ± 0.014 |

a – positive control

b – negative control
